# Targeting the RHOA pathway improves learning and memory in *Kctd13* and 16p11.2 deletion mouse models

**DOI:** 10.1101/2020.05.22.110098

**Authors:** Sandra Martin Lorenzo, Valérie Nalesso, Claire Chevalier, Marie-Christine Birling, Yann Herault

## Abstract

Gene copy number variants (CNV) have an important role in the appearance of neurodevelopmental disorders. Particularly, the deletion of the 16p11.2 locus is associated with autism spectrum disorder, intellectual disability, and several other features. Earlier studies highlighted the implication of *Kctd13* genetic imbalance in the 16p11.2 deletion through the regulation of the RHOA pathway. Here, we target the pathway and rescue the cognitive phenotypes of the 16p11.2 deletion mouse models. We used a chronic administration of fasudil (HA1077), an inhibitor of the Rho-associated protein kinase (ROCK), in mouse models carrying a heterozygous inactivation of *Kctd13*, or the deletion of the entire 16p11.2 BP4-BP5 region. We focused our attention on the most robust cognitive phenotypes seen in the 16p11.2 models and we showed that a chronic fasudil treatment can restore object recognition memory in both mouse models but does not change other behavioural traits. These findings confirm KCTD13 as one target gene causing cognitive deficits in 16p11.2 deletion patients, and the pertinence of the RHOA pathway as a therapeutic path and reinforce the contribution of other gene(s) involved in cognitive defects found in the 16p11.2 CNV models.

**HIGHLIGHTS:**

- Kctd13 haploinsufficiency recapitulates most of the behaviour phenotypes found in the 16p11.2 *Del*/+ models
- Fasudil treatment restores Kctd13 and 16p11.2 *Del*/+ mutant phenotypes in novel location and novel object recognition memory tests
- Fasudil treatment restores the RhoA pathway in *Kctd13*^+/-^ and 16p11.2 *Del*/+ models

## INTRODUCTION

Genetic copy number variants (CNV) of the 16p11.2 locus are an important risk factor for multiple neurodevelopmental disorders (Conrad et al., 2010a; Conrad et al., 2010b; Cook and Scherer, 2008; Nalls et al., 2011; Pinto et al., 2010). The most recurrent 16p11.2 rearrangements, deletion and reciprocal duplication, induce intellectual disability (ID) (Cooper et al., 2011) and Autism Spectrum Disorder (ASD) (Fernandez et al., 2010; Hanson et al., 2015; Jacquemont et al., 2011; Marshall et al., 2008; Weiss et al., 2008; Zufferey et al., 2012). In addition, they are associated with other neuropsychiatric disorders, such as epilepsy (Zufferey et al., 2012), attention deficit / hyperactivity disorder (Shinawi et al., 2010), schizophrenia and bipolar trouble (McCarthy et al., 2009). Body mass index phenotypes and abnormal head size have also been reported in these 16p11.2 rearrangements (Bochukova et al., 2010; Jacquemont et al., 2011; Shinawi et al., 2010; Walters et al., 2010; Zufferey et al., 2012). The most frequent variation of the 16p11.2 region corresponds to change in the genetic interval between *SULT1A1 and SPN1*, encompassing 600 kb and 32 genes. Nevertheless, a study conducted in 2011 found a microdeletion of the 118 kb region, containing MVP, SEZ6L2, CDIPT, ASPHD1 and KCTD13, inside the 16p11.2 genetic interval, segregating with ASD in a family over three generations (Crepel et al., 2011). Thus, this *MVP-KCTD13* region should have a potential key role in the neuropsychiatric features linked to 16p11.2 rearrangement.

The modelling of the 16p11.2 rearrangements through animal models recapitulates the human genetic data. Indeed, three mouse models were developed carrying slightly different deletions of the 16p11.2 homologous genetic interval (Arbogast et al., 2016; Horev et al., 2011; Portmann et al., 2014). Nevertheless, they were found to share common phenotypes including hyperactivity, repetitive behaviours, and deficits in object memory (Arbogast et al., 2016; Horev et al., 2011; Portmann et al., 2014) that are related to human features.

To find the specific brain regions, developmental periods, networks and pathways impacted by the 16p11.2 deletion, several studies have been carried out. In particular, the development of dynamic spatio-temporal networks of 16p11.2 genes integrating data from brain developmental transcriptome with data from physical interactions of 16p11.2 proteins allowed to support the role of KCTD13 as a protein that complex with CULLIN 3 (Cul3) ubiquitin ligase regulating the Ras homolog family member A (RHOA) protein levels (Lin et al., 2015). The known important functions of Rho GTPase signalling pathway in brain morphogenesis at early stages of brain development allowed to propose that *KCTD13* dosage changes in 16p11.2 deletion or duplication carriers influences RHOA levels and lead to impaired brain morphogenesis and cell migration during foetal stages of brain development t(Lin et al., 2015). This hypothesis was supported by later studies showing the implication of KCTD13 in the abnormal brain size associated to 16p11.2 CNVs in zebrafish (Golzio et al., 2012).

In addition, *Kctd13*^+/-^ mouse model showed a reduction of the number of functional synapses, with a decrease of dendritic length, complexity and dendritic spine density due to increased levels of RHOA (Escamilla et al., 2017). These alterations were reversed by RHOA inhibition with rhosin, strengthening the potential role of RHOA as a therapeutic target. Interestingly the recognition deficit was not seen with this model in another more complex paradigm in which the recognition test is based on 3 objects (Escamilla et al., 2017); a more challenging task with multiple step of recognition, involving slightly different mechanisms of memory. Also, recent studies revealed dendritic spine maturation alterations of hippocampal pyramidal neurons in another *Kctd13*^+/-^ mouse model (Arbogast et al., 2019). This model also presented a deficit in recognition and location memory in a paradigm with two objects similar to the phenotype detected in previous studies done on three distinct 16p11.2 *Del*/+ mouse models (Arbogast et al., 2016; Horev et al., 2011; Portmann et al., 2014). Surprisingly, no change in RHOA protein level was detected in the last study (Arbogast et al., 2019). Despite these investigations, how KCTD13 levels can regulate the RHOA signalling pathway, as well as its implication in the phenotypes associated with the 16p11.2 CNVs remain unclear. Thus, we decided to explore the role of *Kctd13* by engineering a new loss-of-function allele, removing only the exons 3 and 4 with CrispR/Cas9, instead of the whole gene (Escamilla et al., 2017) or the exon 2 (Arbogast et al., 2019). Then, we characterized the novel heterozygous mouse model and compared its outcome with the 16p11.2 deletion.

Furthermore, the integration of our results with earlier studies led us to hypothesize that the 16p11.2 deletion leads to an over-activation of the KCTD13-CUL3-dependent RHOA pathway. So, if we assume that the RHOA / ROCK pathway over-activation causes behavioural and learning alterations in the 16p11.2 deletion, inhibiting this pathway should improve the *Kctd13*^+/-^ and the 16p11.2 *Del*/+-associated phenotypes. We decided therefore to treat our mice with fasudil (HA1077), an inhibitor of the ROCK kinase from the RHOA pathway. We used a similar strategy to counterbalance the behavioural impairments of the Oligophrenin-1 mouse model of intellectual disability (Meziane et al., 2016). Thus, we planned first to characterize the behaviour phenotypes of the new *Kctd13* haplo-insufficient model and to compare the outcome with the 16p11.2 *Del*/+ mice. Then we set up a chronic administration of fasudil (HA1077) in adult *Kctd13*^+/-^ and 16p11.2 *Del*/+ mice (this work and (Arbogast et al., 2016)) and evaluated the behaviour of the treated versus the non-treated animals. This evaluation was carried out, focusing on open field activity and object memory (both location and recognition) and molecular analysis of key members of the RHOA pathway.

## MATERIALS AND METHODS

### Mouse lines, genotyping and ethical statement

Two mouse models were used in the study. The 16p11.2 mouse model corresponds to the *Del*(*7Sult1a1-Spn*)6Yah mouse model(Arbogast et al., 2016), noted here *Del*/+. The line was kept on a pure C57BL/6N (B6N) inbred genetic background. The deletion allele was identified by PCR using primers Fwd1 (5’-CCTGTGTGTATTCTCAGCCTCAGGATG-3’) and Rev2 (5’-GGACACACAGGAGAGCTATCCAGGTC-3’) to detect a specific band of 500 bp while the wild-type allele was identified using Fwd1 and Rev1 (5’ - GGACACACAGGAGAGCTATCCAGGTC-3’) primers to detect the presence of a 330 bp fragment. PCR program was: 95 °C / 5 min; 35× (95 °C / 30 s, 65 °C / 30 s, 70 °C / 1 min), 70 °C / 5 min.

The *Kctd13^em2(IMPC)Ics^* knock-out mouse was generated by the CRISPR / Cas9 technology(Birling et al., 2017) in the B6N genetic background. Two pairs of sgRNAs, one pair located upstream and the other pair downstream of the target region, were selected to delete the exon 3 and 4 of the gene. Both pairs of sgRNAs (showing a cut) and Cas9 mRNA were microinjected in fertilized eggs derived from super-ovulated sexually immature B6N female mice (4–5 weeks olds). Injected embryos cultured *in vitro* were implanted into the oviducts of pseudo-pregnant females. The deletion of *Kctd13* in *Kctd13^em2(IMPC)Ics^* (noted here *Kctd13*^+/-^) was confirmed by PCR using primers Ef (5’-ACCTCTTAGCTGGGCATGCTAAATT-3’) and Xr (5’-AGCCTATGCTAACTATTATCACAGG-3’) and the sequence of the deleted fragment. PCR reaction gave deletion and wild-type products of 429 and 668 bp long respectively. PCR program was: 94°C / 5 min, 35 X (94°C / 30 sec; 60°C / 30 sec; 72°C / 30 sec), 72°C / 5 min. This set of primers was also used for genotyping.

Experimental procedures involving animals were approved by the Ministry of National Education, Superior Learning and Research and with the agreement of the local ethical committee Com’Eth (n° 17) under the accreditation number APAFIS#3590-2016011510199843 v4 with YH as the principal investigator (accreditation 67-369).

### Chronic fasudil treatment

In this study, we developed a protocol for a pre-clinical treatment (Figure 1) with the drug fasudil hydrochloride or HA1077 (F4660, LC laboratories Boston, MA, USA). At weaning, control wild type littermates and heterozygous male mice, either *Kctd13*^+/-^ or *Del*/+, were taken from several litters and housed in groups of 4-2 individuals in ventilated cages (Green Line, Techniplast, Italy), where they had free access to water and diet (D04 chow diet, Safe, Augy, France). Animal bedding (Litiere peuplier AB 3 autoclavable, AniBed, Pontvallain, France) was changed once a week. At 11 weeks, animals were transferred from the animal facility to the phenotyping area. The temperature was kept at 21±2 °C, and the light cycle was controlled as 12 h light and 12 h dark (lights on at 7 am).

**Figure 1.**
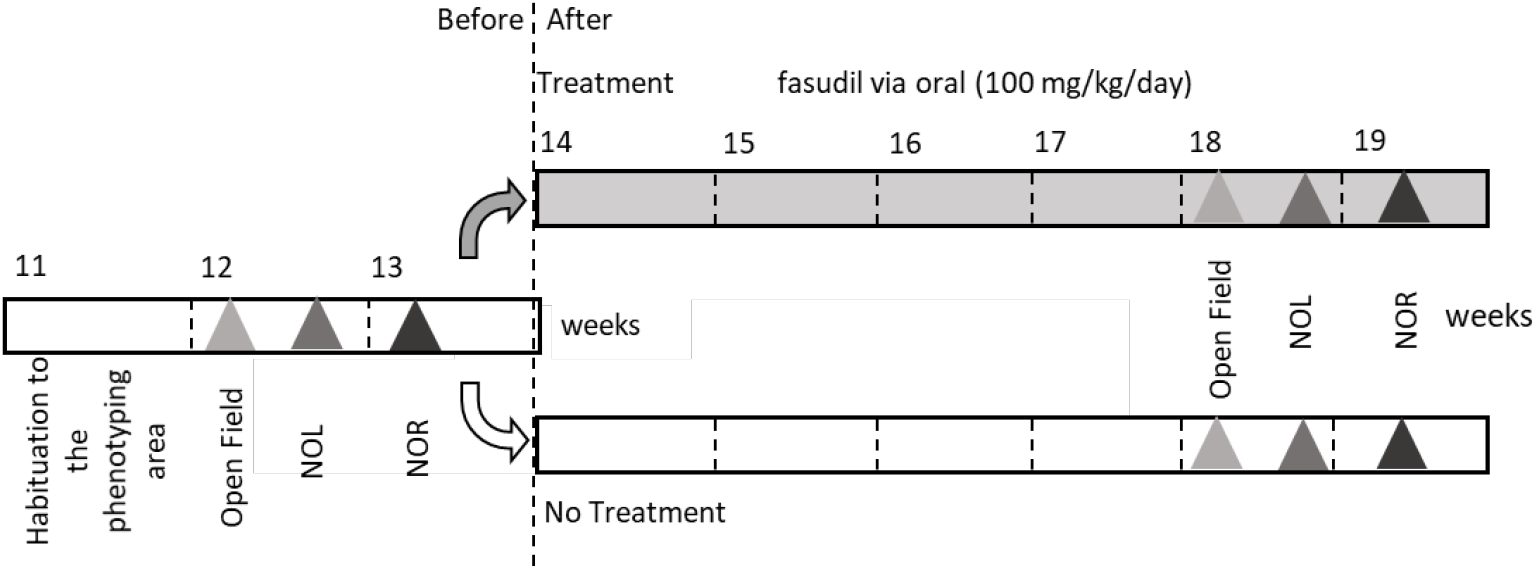
Representation of the behavioral pipeline used for investigating the therapeutic effects of fasudil drug in *Kctd13*^+/-^ and 16p11.2 *Del*/+ mouse models. We used 3 cohorts of animals for each genetic line. 12-week-old mice were subjected to different behavior and learning tests, after a previous habituation to the phenotyping zone. At 14 weeks, each cohort was divided into two groups. The first group started the fasudil treatment and the second group followed the cognitive characterization without treatment. NOL: novel object location, NOR: novel object recognition.

At 12 weeks of age, three independent cohorts of mice for each line, with wild-type (wt) and mutant littermates, were subjected to a battery of behavioural tests (see below) for 2 weeks. Then, 14 weeks old mice were randomly divided into 2 groups: one treated with fasudil administrated orally *ad libitum* in drinking water to reach a dose of 100mg/kg/day and a second with no treatment. The dose was estimated to 152.7 mg in feeding bottle (250 ml) changed twice a week considering the previous study in which we controlled that the drinking volume was about 4.6ml/day/mouse (Meziane et al., 2016). Four weeks after the beginning of the treatment, 18-week-old mice were challenged once again to the same battery of behavioural tests (see below) and were kept under the same treatment condition (Figure 1). The experiments were conducted blindly for genotype as recommended by the ARRIVE guidelines (Karp et al., 2015; Kilkenny et al., 2010). All animals injured by their cage companions were excluded from the behavioural tests at the time when they were seen. A second batch of three independent cohorts were processed similarly but without behaviour test for the molecular analysis of the hippocampal region from treated and non-treated mice. In this case, treated animals started the fasudil treatment at the age of 12 weeks, for 6 weeks. Samples were quickly harvested from 18 weeks-old mice after euthanasia by cervical dislocation and snap frozen for molecular analyses.

### Behavioural analysis

We used three tests that previously unravelled robust phenotypes in the three 16p11.2 mouse models (Arbogast et al., 2016; Horev et al., 2011; Portmann et al., 2014): the open field for the exploration activity, the novel object location and the novel object recognition for the learning and memory in mice.

For the open field (OF) mice were tested in automated arena (44.3 x 44.3 x 16.8 cm) made of PVC with transparent walls and a black floor, and covered with translucent PVC (Panlab, Barcelona, Spain). The arena was divided into central and peripheral regions (8 cm peripheral zone and 28 cm central zone) and homogeneously illuminated at 150 Lux. Each mouse was placed on the periphery of the open field and allowed to explore the apparatus freely for 30 min as a new environment. During each session we analysed the exploration activity by measuring the total distance travelled, evaluated the adaptation of the mice to the environment over time, by splitting the data in intervals of 10 minutes and assessed the vertical activity through the number of rears.

The novel object location (NOL) memory task was carried out in the same open field arena as previously described. NOL stimulate the parahippocampal cortex, the entorhinal cortex and the hippocampus(van Strien et al., 2009). In the first day, mice were habituated to the arena for 30 min at 150 Lux. On the following day, animals went through an acquisition trial during the first 10 min in which they were individually presented to 2 similar objects A. Each object was placed 10 cm away from each one of the corners on the north side of the box. The exploration time of objects A (when the animal’s snout was directed towards the object at a distance ≤1 cm) was recorded. Minimum exploration time was set to 3 s, and mice that did not reach this criterion or did not show any interest for one object were excluded from the study. A 10-min retention trial (second trial) was conducted 5 min later, when one of the familiar objects was displaced to a novel location (B) on the south side and the exploration time (t) of the two objects was recorded for 10 min. We used two identical cylindrical objects of black colour with a white circle on top. In this session, minimum exploration time was set also to 3 s, and mice that did not reach this criterion or did not show any interest (0 s of exploration) for one object were excluded from the study. We verified that no preference was seen during the exploration of the left and right object. The recognition index (RI) was defined as (*t*B / (*t*A + *t*B) × 100). A RI of 50% corresponds to chance level and a significantly higher RI reflects a good recognition of which object was moved in between the two sessions.

The novel object recognition (NOR) memory task is based on the innate preference of rodents to explore novelty involving the perirhinal and entorhinal cortex and the hippocampus. The test was performed in a circular open field of PVC white with opaque walls and floor of 30 cm high and 50 cm diameter. On the first and second days, each mouse was habituated to the arena for 15 minutes at 60 Lux. The following day, we started the NOR sessions. First, each animal was individually given a 10 minutes acquisition trial for the presentation of two identical objects A (either marble or dice) placed at the northeast or northwest of the open field arena. The exploration time of both objects A was recorded. 3 hours later (retention delay in home cages), a 10 minutes retention trial (second trial) was performed. One of the identical object A was replaced with a novel object B at the same position. The exploration time of the two objects (familiar object and novel object) was recorded. The recognition index (RI) was defined as (*t*B / (*t*A + *t*B) × 100). A RI of 50% corresponds to chance level and a significantly higher RI reflects good recognition memory. All mice that did not explore the objects for more than 3 seconds during the acquisition trial or the retention trial or did not show any interest for one object were excluded from the analysis.

### Western blot

Fresh hippocampal tissues were isolated by rapid decapitation/dissection of naive mice and snap frozen. Then, they were lysed in ice-cold sonication buffer supplemented with Complete™ Protease Inhibitor Cocktail (Roche). Individual samples were disaggregated, centrifuged at 4°C for 30 minutes at 14000 rpm, diluted in 4X Laemmli sample buffer containing β-mercaptoethanol (Bio-Rad, France), and incubated at 95 °C for 5 min. Protein concentration was determined by Pierce™ BCA Protein Assay Kit (23225, Thermo Fisher Scientific, Strasbourg). Samples were diluted with sample buffer such that 30 μg of protein were loaded per lane onto 15% polyacrylamide gel. Gels were run and then transferred to nitrocellulose membranes by Trans-Blot^®^ Turbo^™^ Transfer System (BioRad, France) through MIXED MW Bio-Rad Preprogrammed Protocol. Then they were blocked in 5% BSA, 1 X (TBS-T) and incubated with primary antibody during 10 minutes. Membranes were washed in TBS-T followed by a 10 minutes secondary antibody incubation using an HRP conjugated Goat anti-Rabbit IgG (A16096, Invitrogen, France) at 1:5,000 through SNAP i.d.^®^ 2.0 Protein Detection System (C73105, Merck). This apparatus has a vacuum-driven technology and a built-in flow distributor that actively drives reagents through the membrane.

Total levels of RHOA protein and Myosin Light Chain phosphorylation by Myosin Light Chain Kinase via RHOA pathway were analysed using Western Blot. Proteins were visualized with Amersham™ Imager 600. Signals were quantified using ImageJ and analysed using Microsoft Excel and GraphPad Prism. We used the following primary antibodies: RHOA (2117, Cell Signaling, USA, 1:1,000) and pMLC (Thr18/Ser19 #3674, Cell signalling, Boston, MA, USA, 1:1,000). The ratio of protein level or phosphorylation levele against control β-actin protein level (detected with a mouse monoclonal Anti-β-Actin-Peroxidase antibody (A3854 Sigma)) was normalized to untreated wt sample mean.

### Statistical analysis

The statistical analysis was carried out using standard statistical procedures available on the SigmaPlot software (Systat software, San Jose, USA). All outliers were identified using the Grubbs’ test from calculator GraphPad (GraphPad Software, San Diego) or ROUT method with a Q value of 1% from GraphPad Prism 7.01 (GraphPad Software, San Diego) when data with nonlinear regression. Data from behavioural characterization of *Kctd13*^+/-^ and 16p11.2 *Del/+* mouse models were analysed through the Student t-test (see supplementary table 1 and table 2 as a summary). One sample t-test was used to compare recognition index values to the set chance level (50%). Data from post-treatment behavioural phenotyping of both genetic lines were analysed using one-way ANOVA followed by Tukey’s post-hoc test whenever data presented normal distribution and equal variance. Otherwise, we used the non-parametric Kruskal-Wallis one-way analysis of variance and Mann-Whitney *U* test. One sample t-test was used also to compare recognition index values to the set chance level (50%). Western blot data were analysed using Kruskal-Wallis one-way analysis of variance test between groups followed by Mann-Whitney *U* test or Student t-test depending on data distribution. Data are represented as the mean ± SEM and the significant threshold was *p* < 0.05.

## RESULTS

### Phenotypic characterization of the new *Kctd13*^+/-^ mouse model and comparison with the 16p11.2 *Del*/+ mouse model

We created a new *Kctd13* KO mouse model with the deletion of exon 3 and 4, which have been validated by the IMPC (www.mousephenotype.org) and we evaluated its behaviour in three tasks and compared the results obtained with the characterization of 16p11.2 *Del*/+ model (Figure 2). First, we did not detect significant differences in the open field test (12weeks), when we measured the total distance travelled by *Kctd13*^+/-^ mice compared to wt littermates. This result was different from the phenotype of the *Del*/+ mice (figure 2A). Indeed *Del*/+ mutant mice were more active with a significant increase in the distance travelled compared to their wt littermates (Student t test: Total distance: wt vs. *Del*/+ t_(68)_ = −4.096; *p* < 0.001). Then, we analysed the distance travelled in 5-minute intervals, to see the habituation to a new environment during the test. We found that the *Kctd13*^+/-^ mice experience a similar habituation to the control individuals. Here too, a significant difference was seen in the 16p11.2 *Del*/+ carrier mice. Athough being more active, the mutant mice showed normal habituation compared to their wild-type littermates. Finally, we looked at the vertical activity with the numbers of rears, and we did not see any significant differences in the two lines. However, the *Del*/+ mice developed a tendency to the appearance of repetitive behavior, measured from the number of rears with 8 *Del*/+ animals having a strong rearing activity but as a group it was not significant.

**Figure 2.**
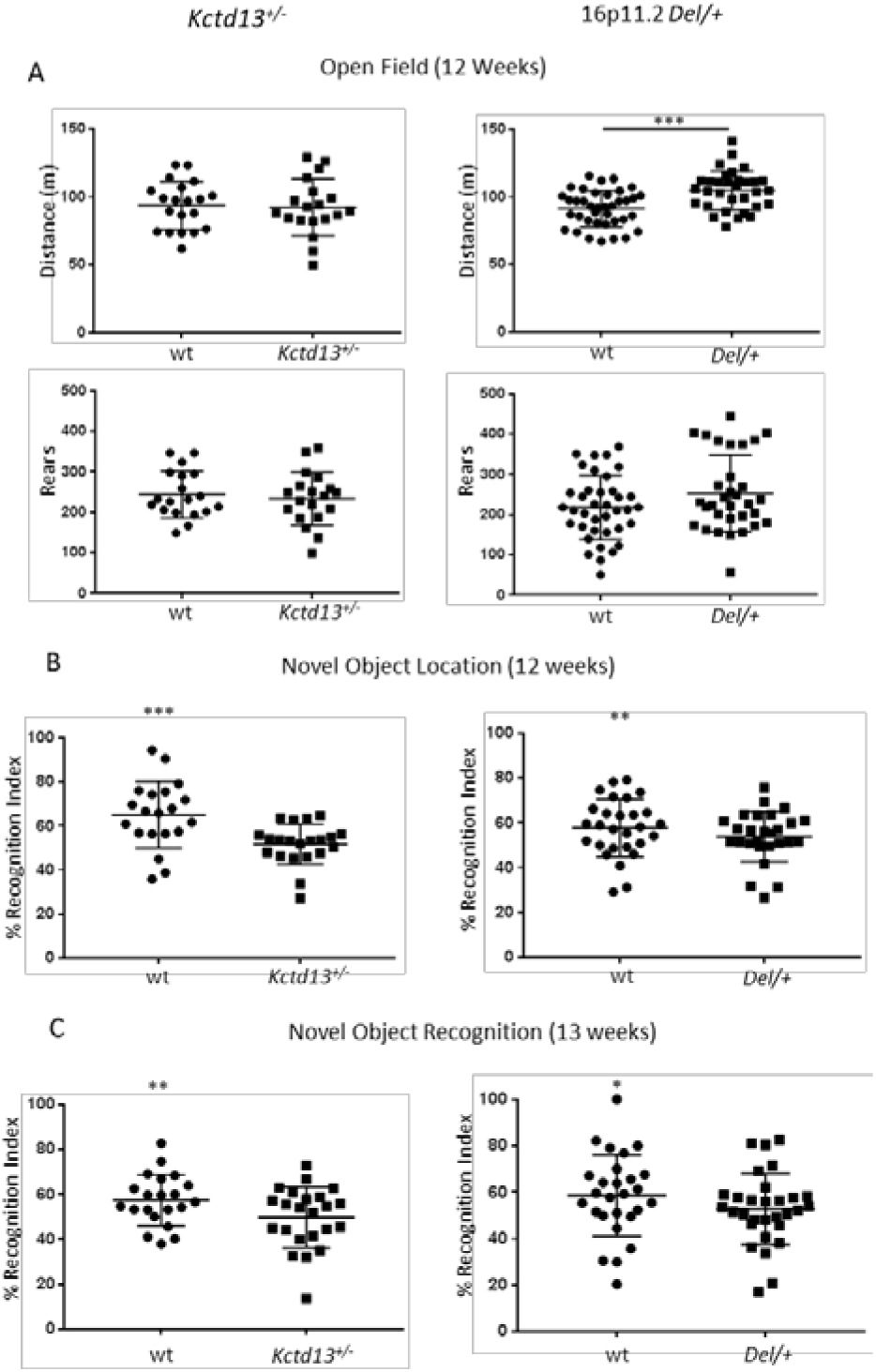
(A) The open field test, (B) the novel object location test and (C) the novel object recognition test of the *Kctd13*^+/-^ and the 16p11.2 *Del*/+ mouse models at 12 and 13 weeks of age. (A) Mutant mice from the *Kctd13*^+/-^ line (wt (n=20) and *Kctd13*^+/-^ (n=19)) showed no alterations in exploratory (total distance) and vertical activity (rears) compared to their wt littermate whereas the mutant mice from the 16p11.2 *Del*/+ line (wt (n=38) and *Del*/+ (n=32)) showed increase exploratory activity in the distance travelled compared to wt. (B) In the NOL test, mutant mice from the two lines (wt (n=20) and *Kctd13*^+/-^ (n=21) littermates; and 16p11.2 *Del*/+ (wt (n=28) and *Del*/+ (n=27) littermates) showed a recognition index not significantly higher to the chance level of 50% and therefore a deficit in object location recognition memory compared to their wt littermates. (C) In the NOR test, the *Kctd13*^+/-^ and the *Del*/+ animals (wt (n=21) and *Kctd13*^+/-^ (n=23) littermates; and wt (n=27) and *Del*/+ (n=30) littermates) showed a poor object recognition memory compared to their respective wt littermates. (* *p* < 0.05; ** *p* < 0.01; *** *p* < 0.001).

Afterwards, we analysed the object location memory (Figure 2B). For this test, animals were scored for their ability to distinguish between a previously presented object that stays in the same location and another object whose position was changed, after a retention delay of 5 min. The recognition indexes for the novel location object showed that, contrary to their wt littermates, both *Kctd13*^+/-^ and 16p11.2 *Del*/+ mice were not able to differentiate between the novel and familiar location. The recognition index for both genotype was not significantly higher to the chance level of 50% (One sample t-test for the *Kctd13*^+/-^ line: wt (t_(19)_ = 4.4607; *p* = 0.0003), *Kctd13*^+/-^ (t_(16)_ = 0.9170; *p* = 0.3728); and the 16p11.2 *Del*/+ model: wt (t_(27)_ = 3.2299; *p* = 0.0032), *Del*/+ (t_(26)_ = 1.8372; *p* = 0.0776)).

Finally, we investigated whether *Kctd13*^+/-^ mice could discriminate a novel object from a previously explored set of two objects after a retention delay of 3 hours in the NOR task (Figure 2C). Whereas wt animals were able to differentiate objects showing a novel object preference, *Kctd13*^+/-^ mice were not able to discriminate the novel from the familiar object. The deficit was like the one seen in the 16p11.2 Del/+ mice (One sample t-test for the *Kctd13*^+/-^ model: wt (t_(20)_ = 3.0179; *p* = 0.0068), *Kctd13*^+/-^ (t_(22)_ = 0.0805; *p* = 0.9366); and the 16p11.2 Del/+ model: wt (t_(26)_ = 2.5312; *p* = 0.0178), *Del*/+ (t_(29)_ = 1.0119; *p* = 0.3199)). Overall, our behavioural analysis showed that the *Kctd13* haploinsufficiency in the pure C57BL/6N genetic background phenocopied the object location and recognition memory deficits seen in the 16p11.2 deletion model. However, the increased exploration activity found in the 16p11.2 *Del/+* mice was not seen in the *Kctd13*^+/-^ mutant mice.

### Fasudil treatment partially reverses the cognitive impairment in the *Kctd13*^+/-^ and in the 16p11.2 *Del*/+ mouse models

Thus, after the behavioural characterization of the *Kctd13*^+/-^ and 16p11.2 *Del*/+ mouse models, we subdivided both genotypes (wt and mutant) into two groups. Individuals were randomly assigned to a group that was treated with fasudil or an untreated control non-treated group prior to further behaviour testing (Figure 1).

First, we noticed that the fasudil treatment did not change the locomotor exploration activity of the different genotypes (Figure 3A) with Del/+ individuals travelling more than the control wt with no effect of the treatment (One way ANOVA between groups: F_(3,63)_ = 10.158; *p* < 0.001; Tukey’s post hoc tests: non treated wt vs. treated wt: *p* = 0,675, non-treated *Del*/+ vs. treated *Del*/+: *p* = 0.54, non-treated wt vs. treated *Del*/+: *p* = 0.001, treated wt vs. non treated *Del*/+: *p* = 0.004 and treated wt vs. treated *Del*/+: *p* < 0.001). The treatment did not have aeffect on any genotype for the vertical activity (Kruskal-Wallis one-way analysis of variance: H(3) = 7.074; *p* = 0.070). Interestingly, one month after the first phenotypic characterization, *Kctd13*^+/-^ mice performed better in the NOL (figure 3B) (One sample t-test: non treated wt (t_(10)_ = 3.9679; *p* = 0.0027), treated wt (t_(11)_ = 5.3506; *p* = 0.0002), non-treated *Kctd13*^+/-^ (t_(12)_ = 2.3628; *p* = 0.0359), treated *Kctd13*^+/-^ (t_(8)_ = 5.7266; *p* = 0.0004)). In contrast, none of this improvement was observed at 18 weeks of age in 16p11.2 *Del*/+ mice (figure 3B) (One sample t-test: non-treated wt (t_(11)_ = 5.4167; *p* = 0.0002), treated wt (t_(13)_ = 4.4492; *p* = 0.0007), non-treated *Del*/+ (t_(7)_ = 0.9837; *p* = 0.3580), treated *Del*/+ (t_(14)_ = 2.1021; *p* = 0.0541)). However, fasudil treatment significantly improved the NOR defect found in both *Kctd13*^+/-^ (One sample t-test: non-treated wt (t_(11)_ = 2.6929; *p* = 0.0209), treated wt (t_(10)_ = 5.7297; *p* = 0.0002), non-treated *Kctd13*^+/-^ (t_(11)_ = 2.6101; *p* = 0.0243 (less than 50%), treated *Kctd13*^+/-^ (t_(9)_ = 3.1937; *p* = 0.0109)) and 16p11.2 *Del*/+ mice ((One sample t-test: non-treated wt (t_(16)_ = 2.3736; *p* = 0.0305), treated wt (t_(8)_ = 2.4481; *p* = 0.0401), non-treated *Del*/+ (t_(12)_ = 0.3787; *p* = 0.7115), treated *Del*/+ (t_(10)_ = 2.9168; *p* = 0.0154)) one week later. This observation confirmed the altered recognition memory in mutants at 19 weeks, detected previously at 12 weeks, and showed the protective effect of fasudil treatment in the *Kctd13*^+/-^ and the 16p11.2 *Del*/+ models (Figure 3C).

**Figure 3.**
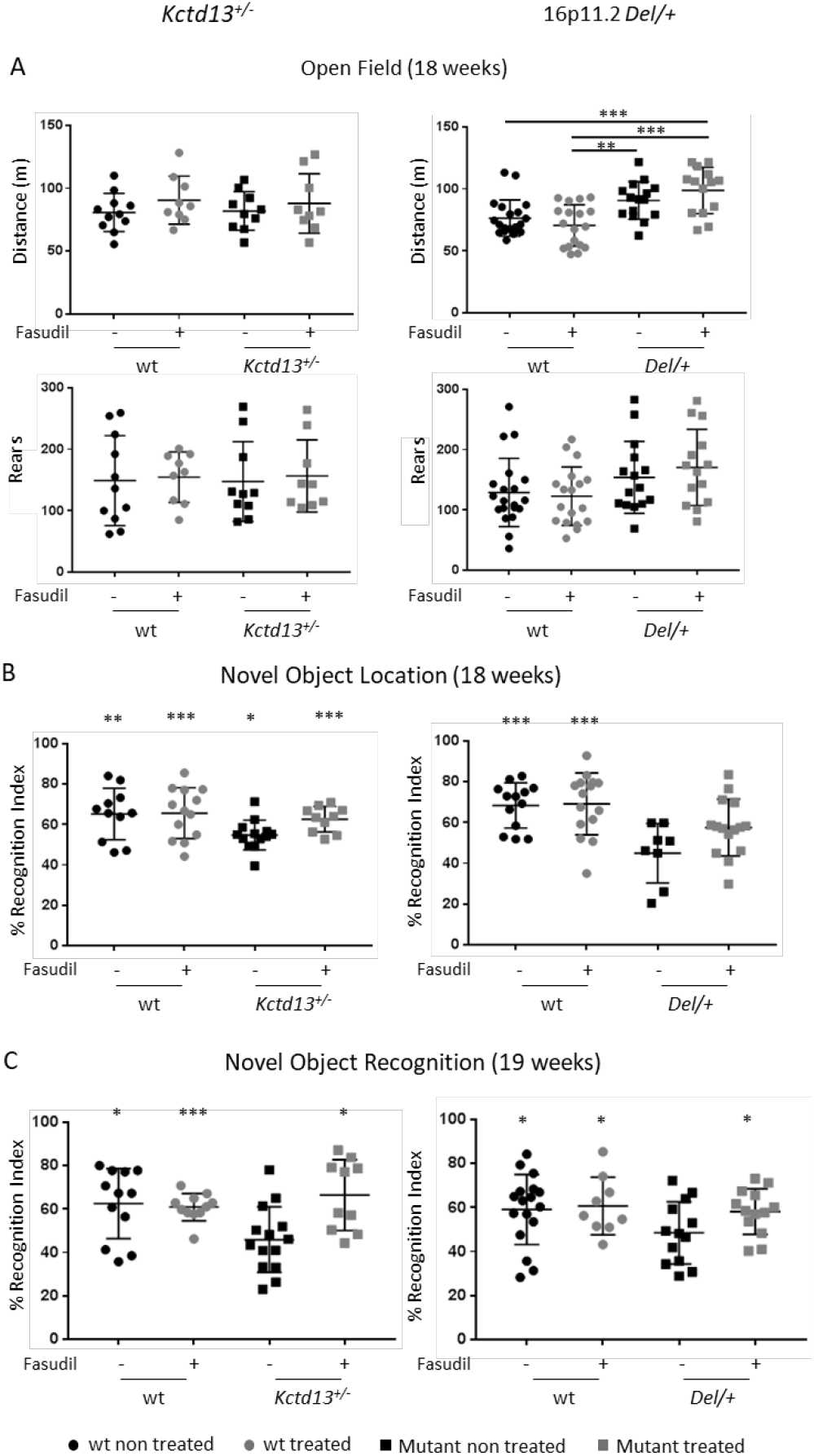
(A) The open field test, (B) the novel object location test and (C) the novel object recognition test of the *Kctd13*^+/-^ and the 16p11.2 *Del*/+ (B) mouse models with or without fasudil treatment. (A) *Kctd13*-deficient mice from the *Kctd13*^+/-^ line (non treated wt (n=11), treated wt (n=9), non-treated *Kctd13*^+/-^ (n=10) and treated *Kctd13*^+/-^ (n=9)) showed no alterations in the different variables compared to their wt littermate. The heterozygous mice from the 16p11.2 *Del*/+ line (non treated wt (n=20), treated wt (n=18), non-treated *Del*/+ (n=15) and treated *Del*/+ (n=14)) showed increased exploratory activity in the distance travelled, which was not affected by the fasudil treatment. The treatment did not have either effect for any genotype on the vertical activity. (B) For NOL test, recognition index reflects the ability of mice from the two lines *Kctd13*^+/-^ (non-treated wt (n=11), treated wt (n=13), non-treated *Kctd13*^+/-^ (n=13) and treated *Kctd13*^+/-^ (n=10) littermates) and the 16p11.2 *Del*/+ (non-treated wt (n=13), treated wt (n=15), non-treated *Del*/+ (n=8) and treated *Del*/+ (n=15) littermates) to distinguish the new location of an object from the familiar one. We observed again that the 16p11.2 *Del*/+ mice showed a deficit in object location memory compared to their wt littermate but the *Kctd13* deficients were no longer defective and the fasudil treatment was not able to restore this ability in the 16p11.2 *Del*/+ model. (C) In the NOR test, the mutant animals from the *Kctd13*^+/-^ (non-treated wt (n=12), treated wt (n=11), non-treated *Kctd13*^+/-^ (n=14) and treated *Kctd13*^+/-^ (n=10)) or the 16p11.2 *Del*/+ model (non-treated wt (n=17), treated wt (n=9), non-treated *Del*/+ (n=13) and treated *Del*/+ (n=13)) were challenged to recognize the new object from the familiar object. *Kctd13*^+/-^ and the *Del*/+ mutant mice were both impaired to recognize the new object compared to their respective wt littermates in the non-treated group and the fasudil treatment was able to restore the object recognition in the *Kctd13*^+/-^ line and in the 16p11.2 *Del*/+ model (* *p* < 0.05; ** *p* < 0.01; *** *p* < 0.001).

### Molecular analyses of RHOA / ROCK signalling pathway in the *Kctd13*^+/-^ and the 16p11.2 *Del*/+ mouse models

Then we checked whether the RHOA / ROCK signalling pathway was changed in both 16p11.2 deletion and *Kctd13* mouse models. Neither the loss of one copy of the complete chromosomic region in 16p11.2 *Del*/+ mice nor the deficiency of *Kctd13* resulted in increased levels of RHOA in the hippocampal region at 18 weeks of age (Figure 4A and B). Nevertheless, both models showed an over activation of the RHOA / ROCK pathway. Indeed, Myosin Light Chain (MLC), a protein targeted by the RHOA/ROCK pathway, showed increased levels of phosphorylation in the hippocampus of these mice, supporting the idea that this pathway could be associated to the cognitive phenotype observed (Figure 4C and D; supplementary table 3). Therefore, we verified whether fasudil therapeutic effect was acting through the inhibition of this RHOA/ROCK signalling pathway. Interestingly, the fasudil treatment reduced MLC phosphorylation levels in *Kctd13* mutant individuals (Kruskal-Wallis one-way analysis of variance between groups: H(3) = 21.731; *p* < 0,001; Mann-Whitney Test: non treated wt vs. non treated *Kctd13*^+/-^: *p* = 0.009; non treated wt vs. treated *Kctd13*^+/-^: *p* = 0.702; non treated *Kctd13*^+/-^ vs. treated *Kctd13*^+/-^: *p* = 0.049). As for the *Kctd13*^+/-^ model, we found that in 16p11.2 deficient mice fasudil restored a normal MLC phosphorylation in treated mutant mice but surprisingly induced increased MLC phosphorylation in wt mice (Kruskal-Wallis one-way analysis of variance between groups: H_(3)_ = 8.457; *p* = 0.037; Mann-Whitney Test: non treated wt vs. treated wt: *p* = 0.008; t-test: non treated wt vs. non treated *Del*/+: *p* = 0.047, non-reated wt vs. treated *Del*/+: *p* = 0.364).

**Figure 4.**
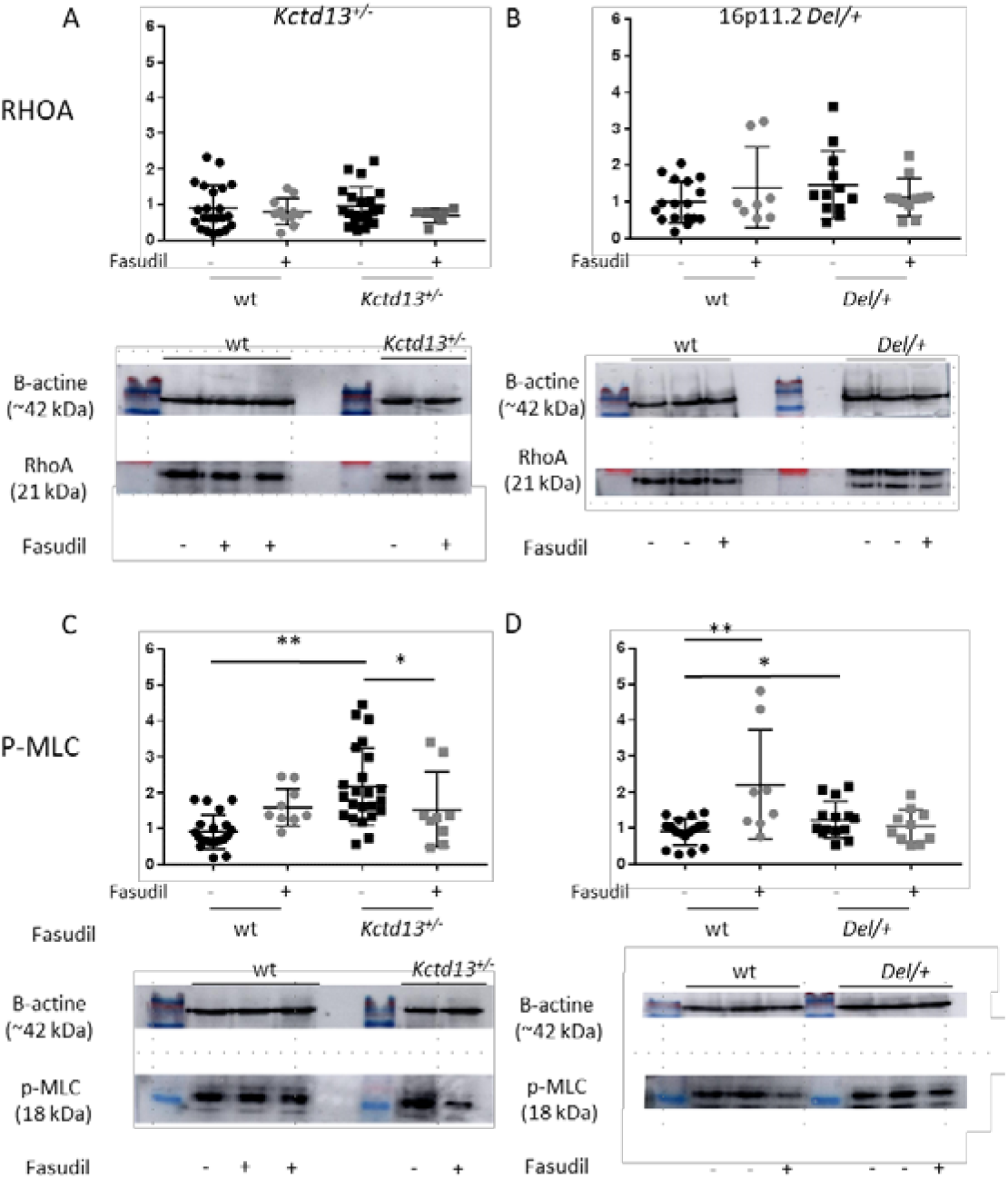
Detection of RHOA (A, B) and of the phoshorylated form of MLC (P-MLC, C,D) by western blots in heterozygous *Kctd13*^+/-^ (A,C) and the 16p11.2 *Del*/+ (B,D) hippocampal lysates and their control (wt) littermate. (A) The quantification of the western blot (an example is shown below the graph) revealed no changes in RHOA protein levels in the *Kctd13*^+/-^ (A) or in the 16p11.2 *Del*/+ (B) mice compared to their wt littermates. Fasudil treatment did not cause changes in RHOA protein levels in the two mutant lines (non-treated wt (n=22), treated wt (n=11), non-treated *Kctd13*^+/-^ (n=21) and treated *Kctd13*^+/-^ (n=7); and non-treated wt (n=17), treated wt (n=8), non-treated *Del*/+ (n=11) and treated *Del*/+ (n=11)). However, *Kctd13* deficient mice showed an increase in the levels of phosphorylated MLC protein (C) and the loss of a copy of 16p11.2 region caused an increase in the levels of phosphorylated MLC protein (D). The treatment with fasudil reversed this alteration in *Kctd13*^+/-^ (non-treated wt (n=21), treated wt (n=10), non-treated *Kctd13*^+/-^ (n=23) and treated *Kctd13*^+/-^ (n=9)) and in the *Del*/+ mutant line (non-treated wt (n=17), treated wt(n=8), non-treated *Del*/+ (n=14) and treated *Del*/+ (n=10)). (* *p* < 0.05; ** *p* < 0.01).

## DISCUSSION

Here, we described the phenotypes of another *Kctd13*^+/-^ mouse mutant line that replicates some of the defects seen in young mouse models carrying the 16p11.2 BP4-BP5 deletion and we explored a treatment aiming at reducing the activity of the RHOA / ROCK pathway on cognition. With the new *Kctd13*^+/-^ mouse line, we were able to detect changes in the NOL and NOR but no effect on the exploration activity compared to the 16p11.2 *Del*/+ mouse model. These results settle with a recent study^(Arbogast et al., 2019)^ highlighting the role of KCTD13 in the 16p11.2 deletion syndromes. The loss of one copy of *Kctd13* gene did not cause alterations on exploration or vertical activity of mice in open field test. However, the hemi-deletion of entire 16p11.2 region induced hyperactivity in these animals. This observation leads us to propose that *Kctd13* genetic dosage is not involved in the increased exploration activity associated with 16p11.2 deletion. Accordingly, chronic fasudil administration did not attenuate the hyper locomotion affecting 16p11.2 *Del*/+ mice. This finding suggests that there could be other genes of the region involved in this phenotype. For this reason, the treatment with an inhibitor of the RHOA / ROCK pathway, deregulated because of *Kctd13* decreased levels in 16p11.2 deficient, did not produce any effect.

When we analysed the NOL in *Kctd13* mutant mice, we found an impairment in the novelty detection in NOL test at 12 weeks of age. However, at 18 weeks when we repeated the test, the non-treated mutant mice showed an improved recognition index. This observation may show that either there is a maturation deficit that is recovered in older mice, may be through compensatory effects, or there could be an effect in the repetition of the test. We favour the first hypothesis as the NOR/NOL phenotypes in this line were found at both ages in the 16p11.2 deletion. Indeed, we found a deep NOL deficit for the 16p11.2 *Del*/+ mice at 12 weeks, that was still present at 18 weeks, not corrected by chronic administration of fasudil for 5 weeks. So, future research will be necessary to analyse how this *Kctd13* NOL phenotype is rescued in 18-week-old no-treated mice. This finding confirms that the NOL-phenotype associated to 16p11.2 rearrangement is not completely dependent on *Kctd13* dosage and its effect on the RHOA / ROCK pathway.

The object recognition memory deficit is one of the most robust and reproducible phenotypes associated with 16p11.2 *Del*/+ mice(Arbogast et al., 2016). In agreement with precedent research, our *Kctd13*^+/-^ mouse model has deficits in novelty detection in NOR. Furthermore, our study showed that fasudil treatment significantly rescued this impairment in mutant mice. Thus, loss of KCTD13 is a driver of object recognition phenotype associated with 16p11.2 deletion. In addition, the therapeutic effects of fasudil on *Kctd13* deficient and 16p11.2 deletion mice highlight the role of the RHOA/ROCK signalling pathway as the main mechanism responsible of this phenotype. Interestingly deficient individuals for the *Kctd13* gene showed no change in expression levels of the RHOA protein and the fasudil treatment did not modify RHOA protein expression in mutant and control mice. Likewise, the carriers of the 16p11.2 hemi-deletion did not display changes of expression for this protein. Even if we were not able to detect a change in the RHOA protein level, *Kctd13* mutants as well as 16p11.2 deficient mice had increased phosphorylated-MLC levels. This observation confirms that RHOA / ROCK pathway outcome is over-activated due to a loss copy of *Kctd13* in *Kctd13* and 16p11.2 mutants. Furthermore, our study showed that the therapeutic effect of fasudil in recognition memory phenotype associated to 16p11.2 CNV was due to the normalizing action of the drug in both mouse models.

At this point, we can highlight the clinical relevance of treatment because of its potential as a cognitive enhancer in humans with memory and learning dysfunction related to neurodevelopmental disorders. However, more work is necessary to understand the precise molecular mechanism affected by the loss of the *Kctd13* gene that is causing the over-activation of the RHOA / ROCK pathway.

We consider that further biochemical studies of the RHOA / ROCK pathway and the KCTD13-CULLIN3 complex are needed to understand the mechanistic role of the pathway in the syndromes associated with 16p11.2 rearrangements. In addition, our results do not prevent that other genes in the region may act at other levels of the RHOA pathway or at different times in development. One of these genes is *Taok2* whose product has been implicated in regulating the activation of the signalling pathway through a functional complex with RHOA, in the cortex of mice exhibiting phenotypes in open field, social preference, fear conditioning, whole brain connectivity, cortical stratification, neuronal morphology, and synaptic function in cortical excitatory neurons(Richter et al., 2019). Probably several genes play a similar role, or the expression of a gene may regulate the expression of another gene in the region, considering the high genotypic density of the 16p11.2 interval and the variability of neurological-associated phenotypes(Girirajan et al., 2012; Jensen and Girirajan, 2019).

## CONCLUSION

Here we showed that *Kctd13* haploinsufficiency phenocopied the object location and recognition memory deficits seen in the 16p11.2 deletion model but did not change the exploration activity in the open field. Treatment in adult mice with fasudil, an inhibitor of RHOA, was able to restore novel object location and novel object recognition memories. Furthermore, fasudil treatment was able to restore RHOA/ROCK signalling pathway to almost normal level of phosphorylation of the Myosin Light Chain in the brain. Altogether targeting the RHOA pathways may be an alternative for improving cognition in people carrying the 16p11.2 deletion.

## ACKNOWLEDGMENT

We are grateful to the animal caretakers for their services at PHENOMIN-ICS and to members of the team, the staff of the IGBMC laboratory for their helpful suggestions and discussions.

## FUNDING

This work has been supported by the National Centre for Scientific Research (CNRS), the French National Institute of Health and Medical Research (INSERM), the University of Strasbourg (Unistra), the French state funds through the “Agence Nationale de la Recherche” under the frame programme Investissements d’Avenir labelled ANR-10-IDEX-0002-02, ANR-10-LABX-0030-INRT, ANR-10-INBS-07 PHENOMIN to YH. The funders had no role in study design, data collection and analysis, decision to publish, or preparation of the manuscript.

## CONFLICT OF INTEREST

The authors have no conflict of interest.

## SUPPLEMENTARY MATERIALS

### SUPPLEMENTARY TABLES

**Supplementary Table 1.** Behavioural characterization of the *Kctd13*^+/-^ mouse model before and after fasudil chronic treatment. In the open field test, no change of horizontal (total distance) or vertical activity (rears) was detected because the inactivation of the gene pre or post treatment. Recognition memory was analysed through the novel object location and object recognition test. *Kctd13*^+/-^ mice showed a deficit for displaced object discrimination and novel object recognition. Mutant males presented no recognition indexes significantly higher than the level of chance 50%. Data are mean ± SEM. Test de Student, * *p* < 0.05, \*\*\**p* < 0.001. One Sample t-test, ^§§^*p* < 0.01, ^§§§^*p* < 0.001 compared with the chance level (50%). Each genotype was divided into 2 groups, non-treated and treated mice. Non-treated *Kctd13*^+/-^ mice showed 4 weeks later improvements for novel object location recognition index. In the novel object recognition test, non-treated *Kctd13*^+/-^ mice showed 4 weeks later a deficit for novel object recognition index while treated *Kctd13*^+/-^ mice recovered the object recognition memory. Data are mean ± SEM. Tukey’s test, * *p* < 0.05. One Sample t-test, ^§^*p* < 0.05, ^§§^*p* < 0.01, ^§§§^*p* < 0.001 compared with the chance level (50%).

| Test | Parameter | Pre treatment |  | Post treatment |  |  |  |
| --- | --- | --- | --- | --- | --- | --- | --- |
|  |  | wt | *Kctd13*<sup>+/-</sup> | wt |  | *Kctd13*<sup>+/-</sup> |  |
|  |  |  |  | Non treated | Treated | Non treated | Treated |
| Open Field | Total distance (m) | 94 ± 4 | 92 ± 5 | 81 ± 5 | 91 ± 6 | 82 ± 5 | 88 ± 8 |
|  | Distance t 0-10 (m) | 40 ± 1 | 38 ± 2 | 34 ± 2 | 35 ± 3 | 33 ± 2 | 34 ± 3 |
|  | Distance t 10-20 (m) | 30 ± 1 | 30 ± 2 | 26 ± 1 | 29 ± 2 | 27 ± 2 | 27 ± 3 |
|  | Distance t 20-30 (m) | 25 ± 2 | 27 ± 2 | 21 ± 2 | 26 ± 2 | 24 ± 2 | 25 ± 3 |
|  | Rears (count) | 244 ± 13 | 234 ± 15 | 149 ± 22 | 155 ± 14 | 148 ± 21 | 157 ± 20 |
| Novel Object Location Recognition<br>5 min delay | S1 object exploration (s) | 9 ± 1 | 10 ± 1 | 7 ± 1 | 6 ± 1 | 8 ± 1 | 7 ± 1 |
|  | S2 non-displaced object exploration (s) | 3 ± 0,4 | 6 ± 1* | 2 ± 0,4 | 3 ± 0,4 | 4 ± 1 | 3 ± 0,4 |
|  | S2 displaced object (s) | 6 ± 1 | 6 ± 1 | 4 ± 0,4 | 6 ± 1 | 5 ± 1 | 6 ± 1 |
|  | Recognition Index (%) | 65 ± 3<sup>§§§</sup> | 52 ± 2<sup>***</sup> | 68 ± 5<sup>§§</sup> | 66 ± 3<sup>§§§</sup> | 55 ± 2<sup>§</sup> | 64 ± 2<sup>§§§</sup> |
| Novel Object Recognition<br>3 hours delay | S1 object A exploration (s) | 18 ± 2 | 19 ± 3 | 13 ± 2 | 30 ± 5* | 19 ± 4 | 16 ± 3 |
|  | S2 object A exploration (s) | 5 ± 1 | 7 ± 1 | 6 ± 1 | 5 ± 1 | 6 ± 1 | 4 ± 1 |
|  | S2 object B exploration (s) | 7 ± 1 | 5 ± 1 | 10 ± 2 | 8 ± 2 | 7 ± 2 | 8 ± 2 |
|  | Recognition Index (%) | 57 ± 2<sup>§§</sup> | 50 ± 3 | 62 ± 4<sup>§</sup> | 61 ± 2<sup>§§§</sup> | 46 ± 4 | 66 ± 5<sup>§</sup> |

**Supplementary Table 2.** Behavioural characterization of the 16p11.2 *Del*/+ mouse model before and after fasudil chronic treatment. In the pre-treatment open field test, the mutant mice showed increased horizontal activity during the 30 minutes of test (total distance) and during the first-time intervals. Recognition memory was analysed in our mouse model through the novel object location and object recognition test. In general, *Del*/+ mice spent more time exploring the objects during session S1. This higher exploration is due to the increased exploration activity associated with the loss of one copy of the 16p11.2 region. Del/+ mice showed a deficit for displaced object discrimination and novel object recognition. Data are mean ± SEM. Test de Student, \*\**p* < 0.01, \*\*\**p* < 0.001. One Sample t-test, ^§^*p* < 0.05, ^§§^*p* < 0.01 compared with the chance level (50%). Our animals were divided into 4 different groups depending on their genotype and treatment. Post-treatment analyses showed for the open field test a higher difference in exploration activity between the treated mutant group and the control group. In the novel object location test, non-treated *Del*/+ animals showed a deficit for displaced object recognition. Treatment in mutant mice increased recognition index but it was not significantly higher than the level of chance (50%). Data are mean ± SEM. Tukey’s test, * *p* < 0.05, \*\**p* < 0.01, \*\*\**p* < 0.001. One Sample t-test, ^§§§^*p* < 0.001 compared with the chance level (50%). In novel object recognition test, non-treated mutant mice showed 4 weeks later a deficit for novel object recognition index while fasudil treatment rescued the object recognition memory in mutant mice. Data are mean ± SEM. Mann-Witney *U* test, * *p* < 0.05. One Sample t-test, ^§^*p* < 0.05 compared with the chance level (50%).

| Test | Parameter | Pre treatment |  | Post treatment |  |  |  |
| --- | --- | --- | --- | --- | --- | --- | --- |
|  |  | wt | 16p11.2<br>*Del/+* | wt |  | 16p11.2<br>*Del/+* |  |
|  |  |  |  | Non<br>treated | Treated | Non<br>treated | Treated |
| Open Field | Total distance (m) | 91 ± 2 | 105 ± 3*** | 76 ± 3 | 71 ± 4 | 91 ± 4 | 99 ± 5*** |
|  | Distance t 0-10 (m) | 35 ± 1 | 42 ± 1 | 30 ± 2 | 29 ± 2 | 34 ± 2 | 35 ± 2 |
|  | Distance t 10-20 (m) | 29 ± 1 | 33 ± 1 | 26 ± 1 | 23 ± 1 | 30 ± 1 | 34 ± 2 |
|  | Distance t 20-30 (m) | 27 ± 1 | 29 ± 1 | 21 ± 1 | 18 ± 1 | 26 ± 2 | 30 ± 2 |
|  | Rears (count) | 217 ± 13 | 253 ± 17 | 129 ± 13 | 123 ± 11 | 154 ± 15 | 171 ± 17 |
| Novel Object<br>Location<br>Recognition<br>5 min delay | S1 object exploration (s) | 10 ± 1 | 16 ± 2** | 7 ± 1 | 9 ± 1 | 12 ± 2 | 10 ± 1 |
|  | S2 non-displaced object<br>exploration (s) | 4 ± 0,4 | 6 ± 1** | 3 ± 0,4 | 2 ± 0,4 | 5 ± 1** | 4 ± 1 |
|  | S2 displaced object (s) | 5 ± 1 | 7 ± 1 | 6 ± 1 | 6 ± 1 | 4 ± 1 | 6 ± 1 |
|  | Recognition Index (%) | 58 ± 2<sup>§§</sup> | 54 ± 2 | 68 ± 3<sup>§§§</sup> | 69 ± 4<sup>§§§</sup> | 45 ± 5** | 58 ± 4 |
| Novel Object<br>Recognition<br>3 hours delay | S1 object A exploration (s) | 16 ± 2 | 20 ± 2 | 15 ± 2 | 11 ± 2 | 30 ± 5* | 24 ± 5 |
|  | S2 object A exploration (s) | 5 ± 1 | 9 ± 1** | 4 ± 1 | 4 ± 1 | 10 ± 2* | 6 ± 1 |
|  | S2 object B exploration (s) | 6 ± 1 | 8 ± 1 | 6 ± 1 | 6 ± 1 | 9 ± 2 | 9 ± 1 |
|  | Recognition Index (%) | 58 ± 3<sup>§</sup> | 52 ± 3 | 59 ± 4<sup>§</sup> | 59 ± 4<sup>§</sup> | 49 ± 4 | 59 ± 3<sup>§</sup> |

**Supplementary Table 3.** RHOA protein and MLC phosphorylation levels analysed by western blot from hippocampal regions of *Kctd13*^+/-^ mouse model treated or non-treated with fasudil Chronic treatment. The intensity of each interest protein bands normalized with the intensity of the corresponding control protein band was again normalized with the mean of all samples of the untreated control individuals. Non-treated *Kctd13*^+/-^ hippocampus presented increased phosphorylation levels of MLC protein while mutant treated with fasudil didn’t present significant increased phosphorylation levels of protein. In the case of the mouse model for the deletion of the 16p11.2 region, we also found an increased phosphorylation levels of MLC. These levels were normalized with the treatment. Mann-Witney *U* test, * *p* < 0.05, ** *p* < 0.01.

| Protein | wt |  | *Kctd13*<sup>+/-</sup> |  |
| --- | --- | --- | --- | --- |
|  | Non treated | Treated | Non treated | Treated |
| RHOA | 0.91 ± 0.14 | 0.81 ± 0.11 | 0.96 ± 0.12 | 0.7 ± 0.07 |
| p-MLC | 0.92 ± 0.1 | 1.6 ± 0.17 | 2.17 ± 0.22** | 1.54 ± 0.35 |

**Supplementary Table 3.** RHOA protein and MLC phosphorylation levels analysed by western blot from hippocampal regions of *Kctd13*<sup>+/-</sup> mouse model treated or non-treated with fasudil Chronic treatment.
| Protein | wt |  | 16p11.2 *Del*/+ |  |
| --- | --- | --- | --- | --- |
|  | Non treated | Treated | Non treated | Treated |
| RHOA | 1 ± 0.14 | 1.4 ± 0.4 | 1.47 ± 0.27 | 1.13 ± 0.16 |
| p-MLC | 1 ± 0.1 | 2.42 ± 0.59** | 1.36 ± 0.15* | 1.16 ± 0.16 |

### SUPPLEMENTARY FIGURES

**Sup figure 1.**
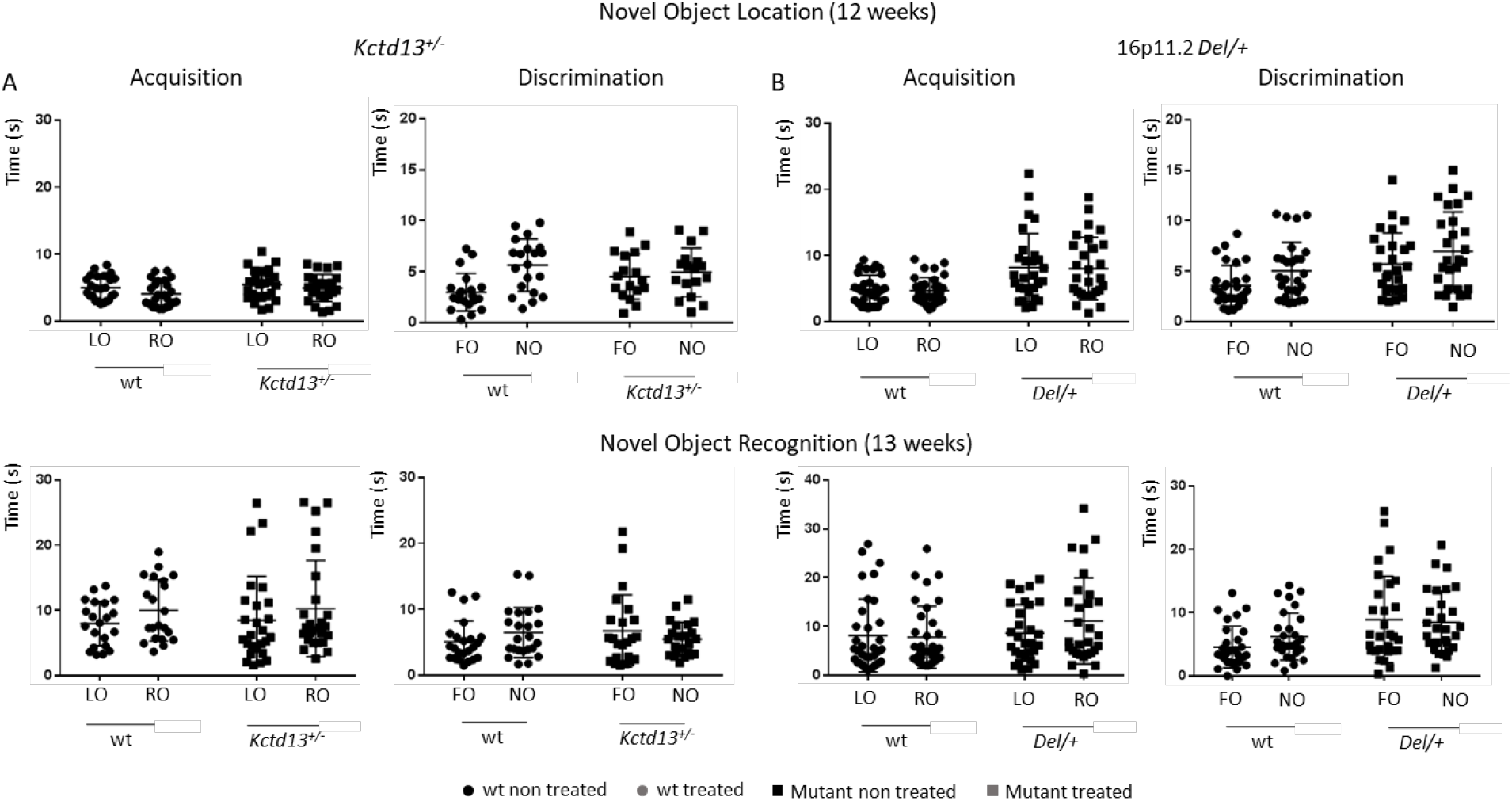
Time spent exploring Novel object location (Top) and novel object recognition (Bottom) in the *Kctd13*^+/-^ (A) and the 16p11.2 *Del*/+ (B) mouse models at 12 and 13 weeks before treatment. No significant change were observed in the absolute exploration time in seconds of the left object (LO) and the right object (RO) in NOL, or familiar (FO) versus novel (NO) object in NOR, during the acquisition trial (10 minutes) in mutant and wt mice.

**Sup figure 2.**
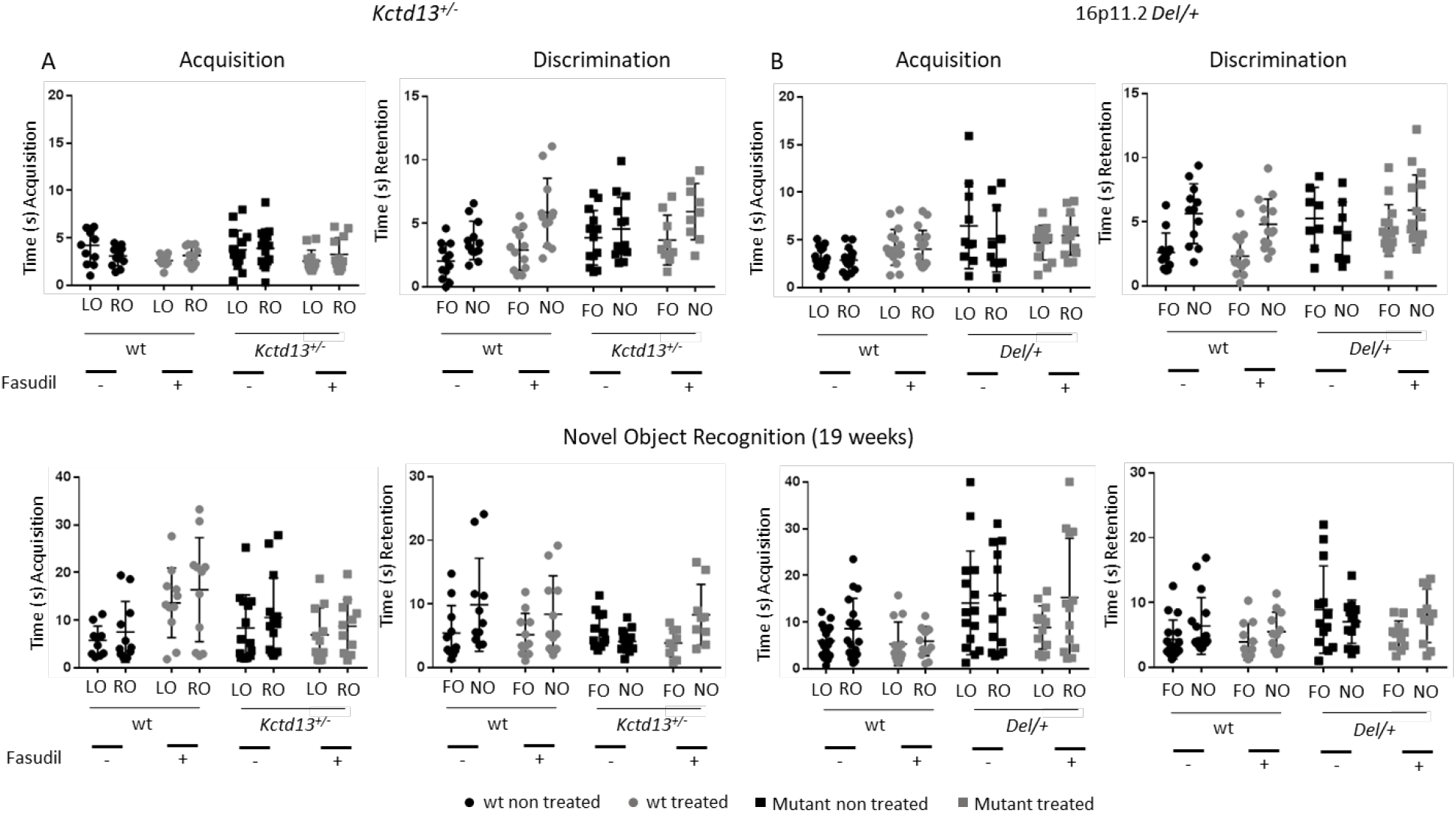
Time spent during Novel object location (Top) and novel object recognition (Bottom) in the *Kctd13*^+/-^ (A) and the 16p11.2 *Del*/+ (B) mouse models at 18 and 19 weeks after treatment. No significant change were observed in the absolute exploration time in seconds of the left object (LO) and the right object (RO) in NOL, or familiar (FO) versus novel (NO) object in NOR, during the acquisition trial (10 minutes) in mutant and wt mice.

**Sup figure 3.**
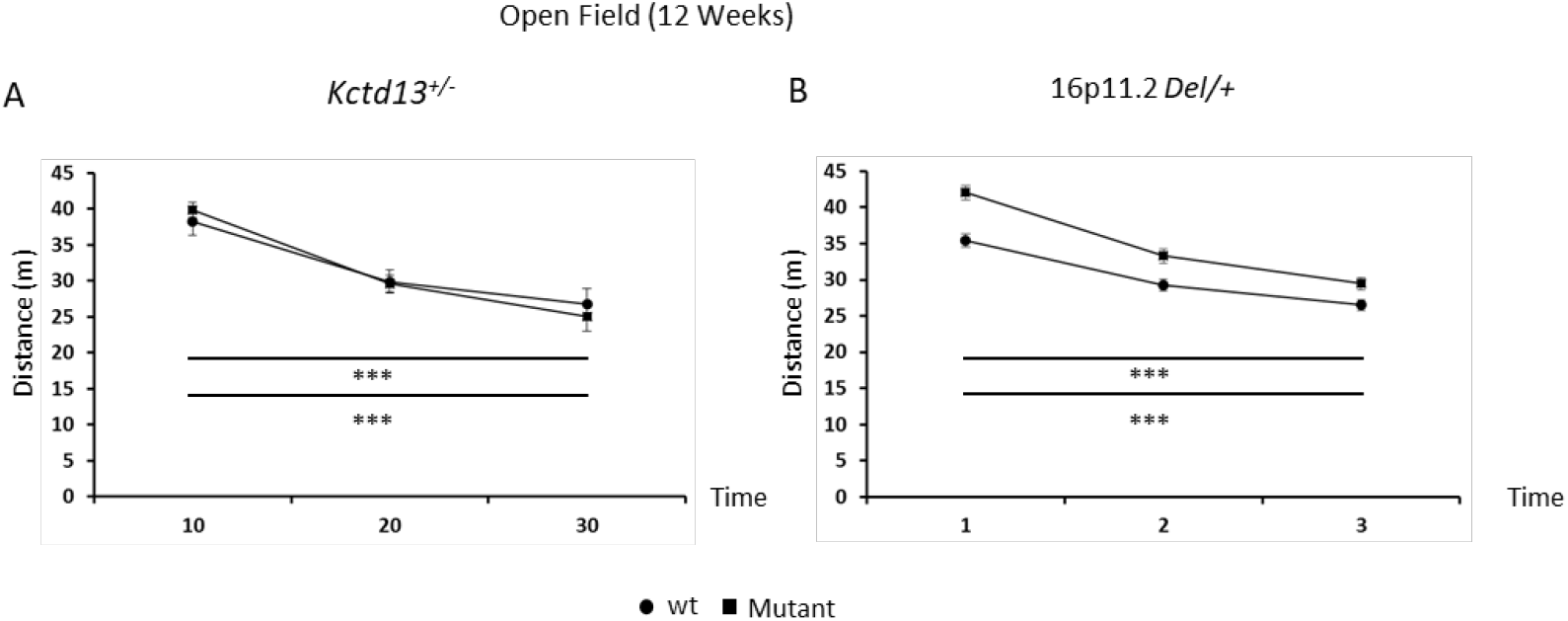
Distance traveled in the open field by the *Kctd13*^+/-^ (A) and *Del*/+ (B) animals showed a normal habituation to the new environment with a significant decreased of the arena exploration during the test. Paired t test: wt: T_0-10_ vs. T_20-30_ t_(19)_ = 7,507; *p* < 0,001; *Kctd13*^+/-^: T_0-10_ vs. T_20-30_ t_(16)_ = 4,453; *p* < 0,001. Paired t test: wt: T_0-10_ vs. T_20-30_ t_(37)_ = 10,834; *p* < 0,001; *Del*/+: T_0-10_ vs. T_20-30_ t_(31)_ = 14,319; *p* < 0,001.

**Sup figure 4.**
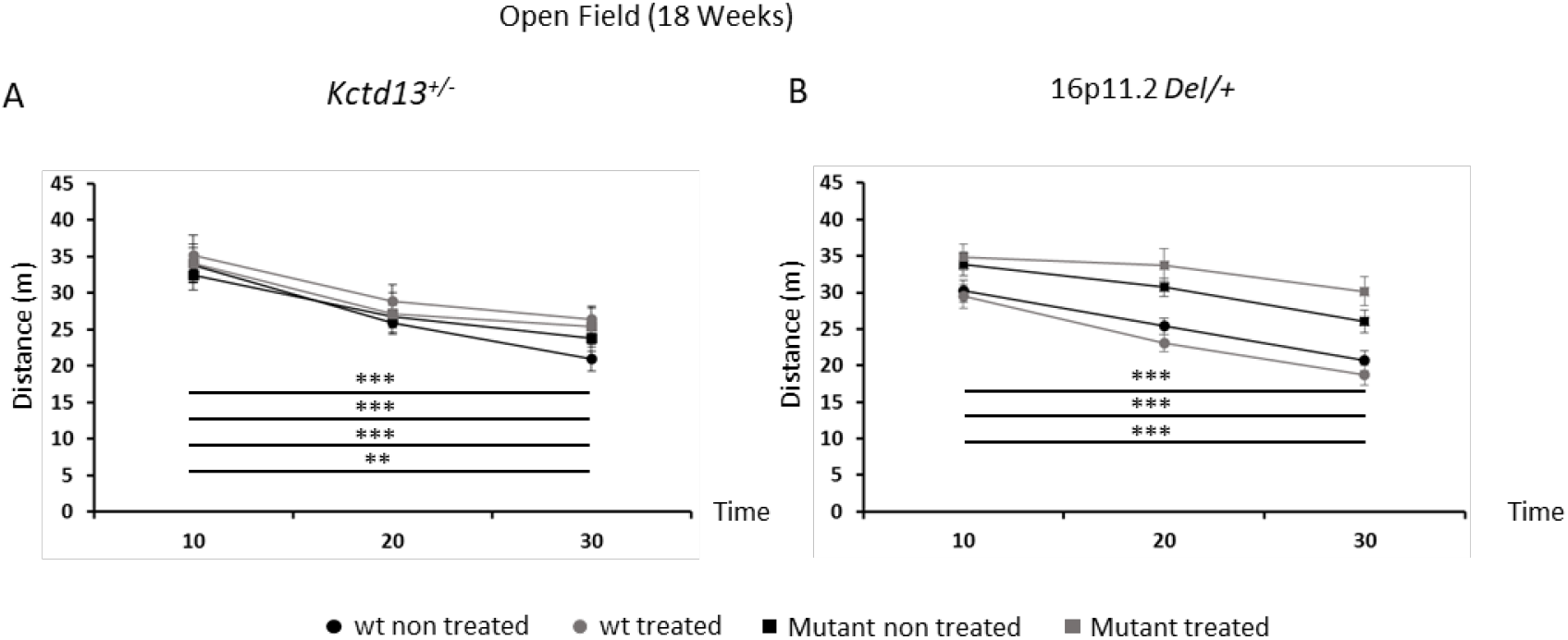
The habituation to a new environment is not altered in the Kctd13 and 16p11.2 deletion models at 18 weeks of ages or by the treatment. The *Kctd13*^+/-^ (A) individuals showed a normal habituation to the new environment with a significant decreased of the arena exploration during the test, which was not affected by the fasudil treatment. Paired t test: non treated wt: T_0-10_ vs. T_20-30_ t_(10)_ = 5,970; *p* < 0,001; treated wt: T_0-10_ vs. T_20-30_ t_(8)_ = 5,117; *p* < 0,001; non treated *Kctd13*^+/-^: T_0-10_ vs. T_20-30_ t_(10)_ = 5,543; *p* < 0,001; treated *Kctd13*^+/-^: T_0-10_ vs. T_20-30_ t_(9)_ = 3,290; *p* = 0.009. Whereas for the *Del*/+ mice (B) fasudil affected the habituation to the environment. Paired t test: non treated wt: T_0-10_ vs. T_20-30_ t_(19)_ = 6,939; *p* < 0,001; treated wt: T_0-10_ vs. T_20-30_ t_(18)_ = 9,753; *p* < 0,001; non treated *Del*/+: T_0-10_ vs. T_20-30_ t_(14)_ = 6,516; *p* < 0,001; treated *Del*/+: T_0-10_ vs. T_20-30_ t_(13)_ = 2,010; *p* = 0.066.

## Notes

### Competing Interest Statement

The authors have declared no competing interest.

